# Protein content and lipid profiling of isolated native autophagosomes

**DOI:** 10.1101/2021.04.16.440117

**Authors:** Daniel Schmitt, Süleyman Bozkurt, Pascale Henning-Domres, Heike Huesmann, Stefan Eimer, Laura Bindila, Georg Tascher, Christian Münch, Christian Behl, Andreas Kern

**Author notes:** **Corresponding author** Andreas Kern, Institute of Pathobiochemistry, The Autophagy Lab, University Medical Center of the Johannes Gutenberg University, 55128 Mainz, Germany.

## Abstract

Autophagy is a central eukaryotic catabolic pathway responsible for clearance and recycling of an extensive portfolio of cargoes, which are packed in vesicles, called autophagosomes, and are delivered to lysosomes for degradation. Besides basal autophagy, which constantly degrades cellular material, the pathway is highly responsive to several stress conditions. However, the exact protein content and phospholipid composition of autophagosomes under changing autophagy conditions remain elusive so far. Here, we introduce a FACS-based approach for isolation of native unmanipulated autophagosomes and ensure the quality of the preparations. Employing quantitative proteomics and phospholipidomics, we obtained a profound cargo and lipid profile of autophagosomes purified upon basal autophagy conditions, nutrient deprivation, and proteasome inhibition. Indeed, starvation only mildly affected the content profile, while interference with proteasome activity showed stronger effects and specifically altered autophagosome cargoes. Interestingly, the phospholipid composition of autophagosomes was unaffected by the different treatments. Thus, the novel isolation method enables purification of intact autophagosomes in large quantities and allows protein content and phospholipid profiling without the requirement of exhaustive cellular fractionation or genetic manipulation.

## Introduction

Macroautophagy (hereafter autophagy) is a eukaryotic catabolic pathway responsible for removal and recycling of an extensive portfolio of cytosolic cargoes. Numerous proteins, aggregates, organelles, cellular compartments, or pathogens have been characterized as autophagy substrates, which are packed in vesicles, called autophagosomes, and are delivered to lysosomes for degradation (Mizushima & Komatsu, 2011).

The engulfment of cargo in vesicles permits its separation and mobilization and is crucial for autophagy-lysosomal degradation. The formation of autophagosomes starts with the double membraned phagophore, which expands around the degradable material until it surrounds it and seals (Lamb *et al*, 2013). This *de novo* vesicle synthesis is initiated by activation of the ULK1/2 kinase complex and the phosphoinositide 3-kinase (PI3-kinase) complex that facilitate the recruitment of proteins and lipids essential for phagophore generation (Mercer *et al*, 2018; Nascimbeni *et al*, 2017). Upon nucleation from the ER membrane, the phagophore elongates by addition of lipids. While the exact lipid sources remain to be elucidated, recent studies suggest a direct lipid transfer from the ER via ATG2A/B (Osawa *et al*, 2019; Valverde *et al*, 2019). Indeed, in yeast it has been shown that phospholipid (PL) synthesis from free fatty acids is important for efficient autophagosome formation (Schutter *et al*, 2020).

The elongation of the phagophore is linked to two ubiquitin-like conjugation reactions (Mizushima, 2020). Firstly, ATG12 is conjugated to ATG5, which secondly facilitates the conjugation of Atg8 proteins to the phospholipid phosphatidylethanolamine (PE) (Kabeya *et al*, 2004; Noda & Inagaki, 2015). In humans, six Atg8 family members have been identified: MAP1LC3A, MAP1LC3B, MAP1LC3C (shortly LC3A-C), as well as GABARAP, GABARAPL1, and GABARAPL2 (Slobodkin & Elazar, 2013). Lipidated Atg8 proteins are inserted into both sides of the growing phagophore double membrane and stay attached to mature autophagosomes, facilitating phagophore elongation and closure as well as autophagosome-lysosome fusion (Nguyen *et al*, 2016; Tsuboyama *et al*, 2016; Weidberg *et al*, 2010).

Additionally, Atg8 proteins provide binding sites for cargo receptors. Autophagy was initially described as a non-selective process that degrades random material of the cytosol to recycle building blocks to respond to changing metabolic requirements. The identification of cargo receptors, though, established a selective part by which specific substrates are directed into the pathway (Khaminets *et al*, 2016; Stolz *et al*, 2014). Via its LC3 interacting domain and its ubiquitin binding domain, the cargo receptor p62/SQSTM1 e.g. binds to LC3 and mediates the transfer of selective cargoes into autophagosomes (Gatica *et al*, 2018; Kirkin *et al*, 2009; Pankiv *et al*, 2007). The formation of autophagosomes is highly dynamic and rapidly adapts to changing cellular conditions. Healthy, unstressed cells are characterized by a basal autophagy rate that constantly degrades cellular material at (comparably) low levels. Various stress situations, such as starvation or imbalances of proteostasis, alter autophagy, resulting in a substantially enhanced cargo degradation (He & Klionsky, 2009). However, the exact profile of the cellular material that is degraded upon different autophagy conditions is not defined so far. Several studies have investigated autophagy substrates upon autophagosome isolation by elaborate cellular fractionation methods with clear limitations in purity (Dengjel *et al*, 2012; Gao *et al*, 2010; Mancias *et al*, 2014). More recently, autophagosome content was examined by proximity labelling in combination with quantitative proteomics, which requires genetic manipulation of the cell (Le Guerroue *et al*, 2017; Zellner *et al*, 2021). Thus, extensive comparative cargo and phospholipid analyses of unmanipulated native human autophagosomes from basal and altered autophagy conditions are missing so far.

Here, we now introduce a FACS (fluorescence activated cell sorting)-based isolation approach to purify large quantities of intact native autophagosomes. We characterized the quality of the isolated autophagic vesicles and conducted quantitative proteomic and phospholipidomic analyses to generate profound autophagosome content profiles and PL compositions upon basal autophagy conditions, nutrient deprivation, and proteasome inhibition. Importantly, starvation only mildly affected the autophagosome cargo content profile, while disturbances of proteasomal degradation showed treatment-specific alterations. The PL composition of autophagosomes was unaffected by the different autophagy modulations, emphasizing the requirement for a distinct PL pattern in autophagic vesicles.

## Results and Discussion

### Isolation of intact native autophagosomes in large quantities

We established a protocol for isolation of native unmanipulated autophagosomes, employing antibody-based fluorescence tagging of Atg8 proteins and subsequent sorting via FACS (Figure 1A, Appendix Figure S1A). Lipidated Atg8 proteins are inserted into the outer as well as inner membrane of elongating phagophores and stay attached to mature autophagosomes (Kabeya *et al*., 2004). Upon mild cell disruption, we incubated the cellular extract with a primary antibody directed against LC3B or GABARAP, followed by treatment with a secondary fluorophore-conjugated antibody. The stable and selective attachment of the fluorophore allowed the specific purification using FACS. The sorting resulted in preparations of approx. 1.000 autophagosomes per µl PBS and thus allowed their purification in large quantities.

**Figure 1.**
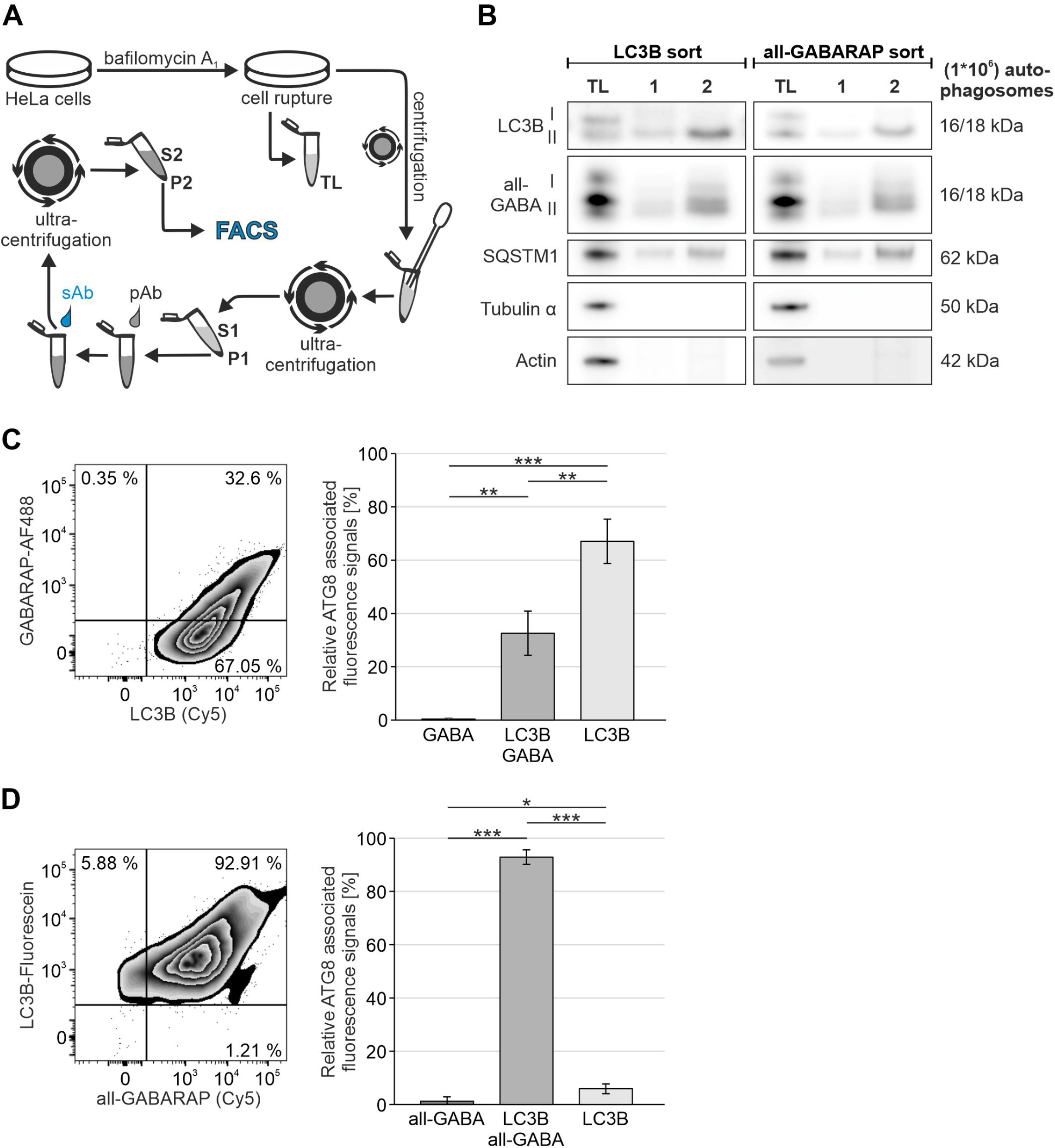
FACS-based isolation of autophagosomes. A. Schematic representation of the FACS-based autophagosome isolation method. TL total lysate; P pellet fraction; S supernatant. B. Western blotting of different numbers of purified autophagosomes. Isolations were based on LC3B or all GABARAP antibodies, respectively, and are represented with total lysate (TL). Depicted are representative blots of 12 independent approaches. C. Co-localization of fluorescence signals linked to LC3B and GABARAP. Shown percentages represent the average distribution of 3 independent experiments excluding double negative events. D. Co-localization of fluorescence signals linked to LC3B and all GABARAP isoforms. Shown percentages represent the average distribution of 3 independent experiments excluding double negative events. C+D. Statistics are depicted as mean ± SD; One-Way ANOVA; *p ≤ 0.05; **p ≤ 0.01; ***p ≤ 0.001.

In order to confirm the quality of the isolation, we analyzed the presence of autophagosome proteins within the isolate fraction by methanol/chloroform-based protein precipitation and Western blotting. Upon preparation of autophagic vesicles via antibodies directed against LC3B or all GABARAP isoforms (Figure 1B), we detected the lipidated variants of both Atg8 proteins as well as p62/SQSTM1 within the isolate fractions. But unlipidated Atg8 proteins, which are not bound to autophagosomes, and cytoskeleton proteins, were hardly detectable. They were quantitatively excluded by the centrifugation steps integrated in the protocol and the fluorescence-based sorting (Figure 1A, Appendix Figure S1B).

To examine the specificity of the antibody binding and the quality of the preparations, we quantified the co-localization of antibody-mediated fluorescence signals of two different autophagosome proteins. We employed antibodies directed against LC3B, GABARAP, or all GABARAP isoforms and analyzed the fluorescence signal via FACS (Figure 1C+D, Appendix Figure S2). Indeed, almost 100 % of GABARAP-positive and all GABARAP-positive structures co-localized with the LC3B antibody-mediated fluorescence signal. Focusing on LC3B, the GABARAP signal was found on approx. 33 % and the all GABARAP signal on around 93 % of LC3B-marked autophagosomes, respectively. This indicated that the different GABARAP isoforms not necessarily co-localize on single autophagosomes. However, the vast majority of vesicles was indeed positive for two different autophagosome markers, emphasizing the exceptional quality of the isolation approach. Only a minor fraction of autophagosomes showed exclusively one Atg8 family member, LC3B (∼ 6 %) or at least one of the GABARAP isoforms (∼ 1 %). This can be explained in different ways: (1) insufficient antibody binding; (2) a small fraction of autophagosomes is indeed only decorated by one particular Atg8 family member; (3) besides the association with autophagosomes, LC3B is also linked to non-autophagic vesicles. During LC3-associated phagocytosis LC3B-II is inserted into the single membrane of phagosomes, supporting lysosomal degradation of phagosome content (Sanjuan *et al*, 2007). However, we treated cells with bafilomycin A_1_ prior to autophagosome preparation, which causes a massive accumulation of autophagosomes within the cell. Thus, LC3-associated phagosomes should only represent a minor fraction of vesicles devoid of GABARAP isoforms.

Continuing our quality control, we investigated whether the purified autophagosomes were sealed and used proteinase K digestion to analyze its impact on p62/SQSTM1 (Figure 2A). In intact vesicles, the cargo receptor is inaccessible and thus protected from degradation (Velikkakath *et al*, 2012). Indeed, p62/SQSTM1 was not prominently degraded by proteinase K within the isolate fractions, indicating that the overall majority of autophagosomes was closed. For control, we opened autophagic vesicles mechanically, which resulted in the exhaustive digestion of p62/SQSTM1.

**Figure 2.**
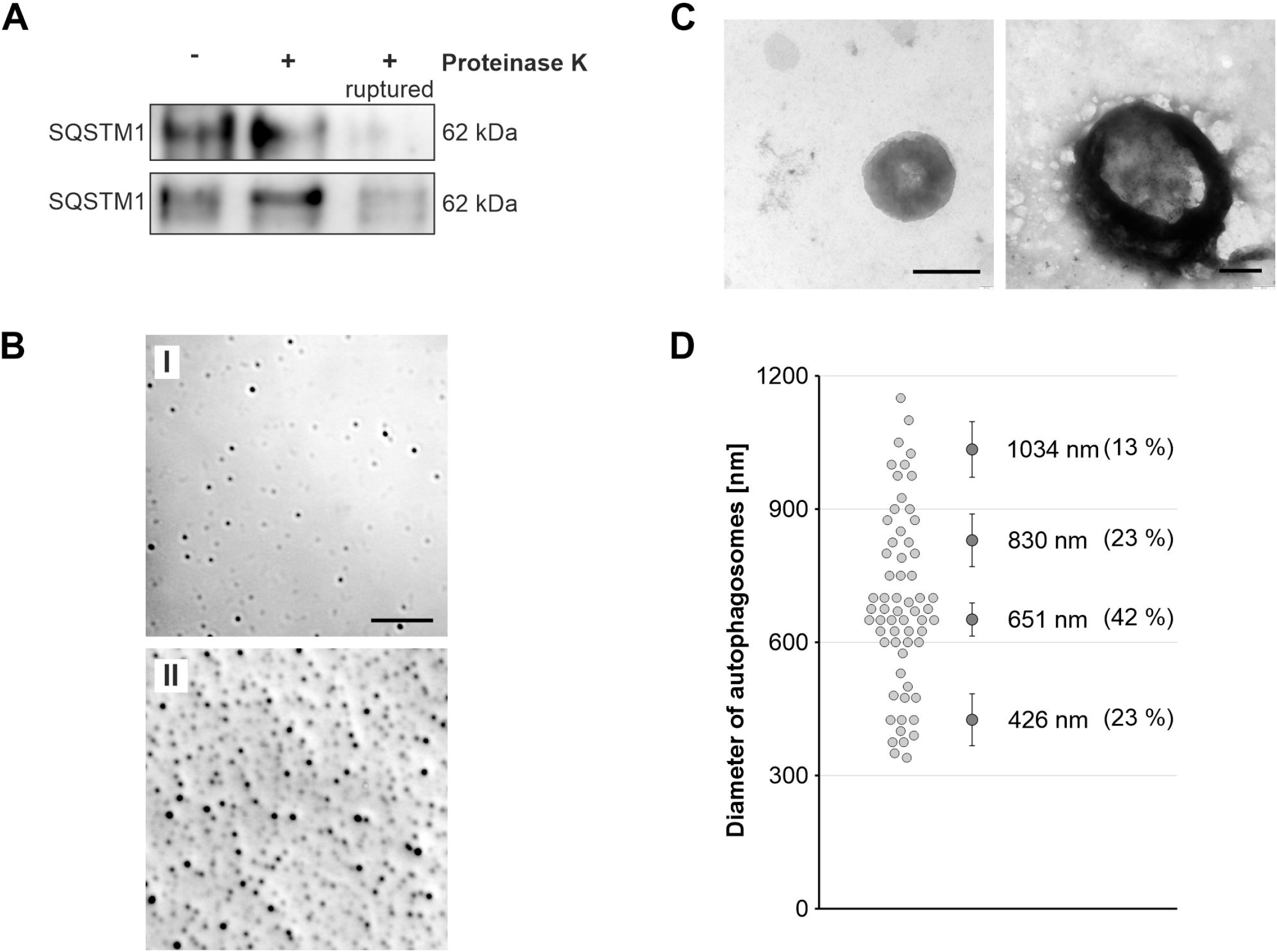
FACS-based isolated autophagosomes are sealed. A. Western blotting of isolated autophagosomes upon proteinase K digestion. Mechanically opened vesicles served as positive control. For negative control, isolates were incubated with BSA instead of proteinase K. Depicted are two different blots that are representative for 5 independent experiments. B. DIC microscopy images of purified autophagosomes at high (I) or low (II) dilution. Images are representative for 3 independent approaches. Scale bar = 10 µm. C. Negative stain electron microscopy images of isolated autophagosomes. Scale bar = 500 nm. D. Size evaluation of isolated autophagosomes. The diameter of approx. 60 individual autophagosomes was determined using EM images. Statistics are depicted as mean ± SD.

Further analyses of the autophagosome preparations were conducted by microscopy. Differential interference contrast (DIC) microscopy showed vesicular structures of different sizes without evidence of cellular debris or accumulations of other cellular material (Figure 2B). This finding was validated by negative stain electron microscopy, which identified double membraned vesicles (Figure 2C). The inner core often showed a granular appearance and was filled with darkly stained structures most likely resembling proteinous cargo. Thus, visualization of the isolated vesicles also confirmed the preparation of pure and undamaged autophagosomes.

Autophagic vesicles are described to cover size ranges of 500 to 1.500 nm in diameter (Mizushima *et al*, 2002). Size evaluation of the purified autophagosomes by EM demonstrated that they showed expected diameter ranges from 340 nm to 1.150 nm (Figure 2D). Interestingly, the autophagic vesicles could be clustered into different size groups. The group with the smallest diameters covered 426 nm on average, the largest grouped with a mean diameter of 1.034 nm. Covering 42 % of all analyzed vesicles, the most abundant autophagosomes showed a mean size of 651 nm in diameter.

### Protein cargo profiles of native autophagosomes upon basal autophagy conditions, starvation, and proteasome inhibition

The established FACS-based protocol enabled the isolation of pure unmanipulated autophagosomes at high quantities. The vesicles were not damaged by the preparation, which qualified the isolates for bulk cargo analyses. For protein content profiling, we conducted quantitative LC-MS/MS and analyzed purified autophagosomes upon basal autophagy conditions (full medium), EBSS-mediated starvation, and MG132 treatment.

EBSS treatment causes a rapid increase in autophagic activity to recycle building blocks to compensate the lack of nutrients (He & Klionsky, 2009). Autophagy is regulated by the MTORC1 complex (Condon & Sabatini, 2019), which, in its active form, facilitates anabolic pathways and restricts autophagic activity. Upon starvation, MTORC1 is inactivated, resulting in the promotion of catabolic processes and in autophagy induction.

MG132 inhibits the proteasome and triggers an imbalance of proteostasis, causing the accumulation of ubiquitinated proteins within the cell. The proteasome is the degradative active component of the ubiquitin proteasome system (UPS), which, in addition to autophagy, is a prominent cellular degradation pathway (Pohl & Dikic, 2019). Importantly, autophagy and the UPS are functionally interrelated. Perturbation of the UPS results in the enhanced autophagic degradation of proteasome substrates, which is regulated by the co-chaperone BAG3 (Gamerdinger *et al*, 2009).

To confirm the impact of EBSS and MG132 on autophagy, we conducted Western blotting and immunocytochemical analyses (Appendix Figure S3). As expected, EBSS treatment increased autophagic activity, since the flux of LC3B-II was enhanced (Appendix Figure S3A). This was validated by immunocytochemical stainings of LC3B-positive autophagosomes, which accumulated at high levels in comparison to control cells when lysosomal degradation was inhibited by bafilomycin A_1_ (Appendix Figure S3B). For proteasome inhibition, we treated cells for 8 h with MG132 and added bafilomycin A_1_ in the last two hours. This treatment resulted in a reduced LC3B-II flux and a decreased accumulation of LC3B-positive structures in comparison to control cells (Appendix Figure S3). The duration of proteasome inhibition and the resulting disturbance of proteostasis apparently exhausted the autophagic system. In summary, both treatments modulated autophagy and thus potentially altered the cargo profile compared to basal autophagosomes.

The proteomics analysis of basal autophagosomes detected 4.514 cargo proteins in total, including multiple autophagosome proteins (Source data, Appendix Figure S4). Functional clustering of the top 500 cargo proteins demonstrated that basal autophagic protein substrates grouped into prominent cellular pathways, such as protein and RNA metabolism or vesicle trafficking (Figure 3A). Comparative analysis of the three independently performed autophagosome preparations upon basal conditions showed only subtle changes (Appendix Figure S5A), confirming the quality and reliability of each independent autophagosome preparation, protein processing, and quantitative MS.

**Figure 3.**
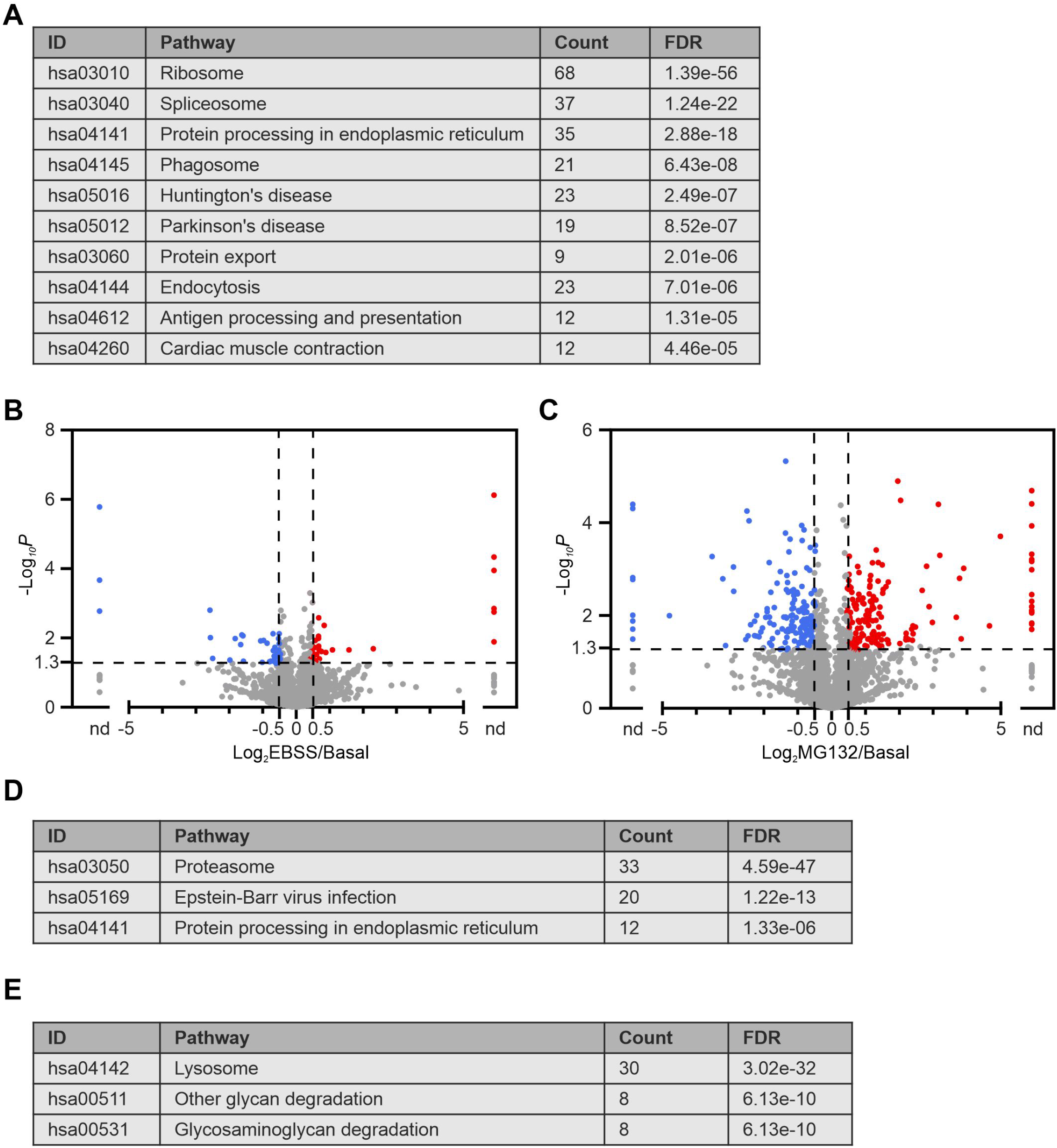
Protein cargo profiles of isolated autophagosomes upon different autophagy conditions. A. KEGG pathway analysis of top 500 basal cargo proteins. Pathways are shown with the number of proteins found in the data set and computed FDRs for enrichment. Represented are the top 10 pathways. B. Volcano plot showing the differential appearance of proteins in autophagosomes of EBSS treated cells in comparison to control cells kept in full medium. Log_2_-transformed fold changes. For proteins that were excluded from autophagosomes upon EBSS treatment or which exclusively appeared within these vesicles, no fold changes could be calculated and are indicated as not determinable (nd). C. Volcano plot showing the differential appearance of proteins in autophagosomes of MG132 treated cells in comparison to control cells kept in full medium. Log_2_-transformed fold changes. For proteins that were excluded from autophagosomes upon MG132 treatment or which exclusively appeared within these vesicles, no fold changes could be calculated and are indicated as not determinable (nd). D. KEGG pathway analysis of proteins with enhanced autophagosome levels upon MG132 treatment. Pathways are presented with the number of proteins found in in the data set and computed FDRs for enrichment. E. KEGG pathway analysis of proteins with reduced autophagosome levels upon MG132 treatment. Pathways are presented with the number of proteins found in in the data set and computed FDRs for enrichment. Represented are the top 3 pathways. See Appendix Figure S7 for complete list.

Investigating autophagosome content upon EBSS treatment detected only minor alterations despite a significant increase in autophagic flux (Figure 3B, Appendix Figure S5B+C). Interestingly, a subset of proteins with increased autophagosome appearance clustered within the ribosome biogenesis pathway (Appendix Figure S5B), which might reflect the reduction of cellular anabolic activities mediated by inhibition of MTORC1. However, the minor influence of nutrient deprivation on the autophagosome protein content profile emphasizes that this condition results in a globally increased turnover of autophagy substrates without selection of particular cargoes. The requirement to rapidly compensate the lack of nutrients apparently restricts the choice of dispensable cellular material.

In contrast to starvation, proteasome inhibition showed a much stronger impact on autophagosome content, resulting in an altered appearance of multiple proteins (Figure 3C). Importantly, network analysis of proteins with an increased autophagosome appearance revealed that a subset grouped within the proteasome pathway (Figure 3D, Appendix Figure S6A), illustrating an enhanced degradation of UPS components upon MG132 treatment. These included multiple proteasome subunits as well as proteins of the ubiquitination process and interestingly also BAG3. Upon proteasome inhibition, BAG3 up-regulates autophagy and mediates the enhanced transfer of ubiquitinated cargoes into autophagosomes (Arndt *et al*, 2010; Gamerdinger *et al*., 2009). Contrary to the enhanced degradation of UPS components, proteins with reduced autophagosome appearance upon MG132 treatment clustered within the lysosome pathway (Figure 3E, Appendix Figure S6B, Appendix Figure S7), including LAMP proteins and several acidic hydrolases, such as cathepsins or glycosidases. Thus, inhibition of the proteasome altered the autophagosome cargo profile and the observed changes displayed an adequate adaptation to the treatment: components supporting UPS function were degraded at enhanced levels, whereas factors supporting the lysosome showed a reduced turnover.

In conclusion, analysis of autophagosome content resulted in a detailed representation of autophagy cargoes and showed treatment-specific effects. This emphasizes the quality of the autophagosome preparations and highlights the value of the isolation to investigate content profiles from native unmanipulated autophagic vesicles. The data depicts a profound basis for the definition of autophagic substrates under various autophagy conditions.

### Phospholipid composition of native autophagosomes

In addition isolation of autophagic vesicles enabled the detailed investigation of the autophagosome PL composition. Employing targeted multiplex quantitative MS, we quantitatively determined different PL species with diverse fatty acid combinations, concerning chain length and saturation level (Figure 4). Importantly, the PL were unaffected by the different autophagy conditions, despite clear alterations in the rate of autophagosome formation (Appendix Figure S3). This indicates the importance of a distinct PL composition to support autophagosome structure and function.

**Figure 4.**
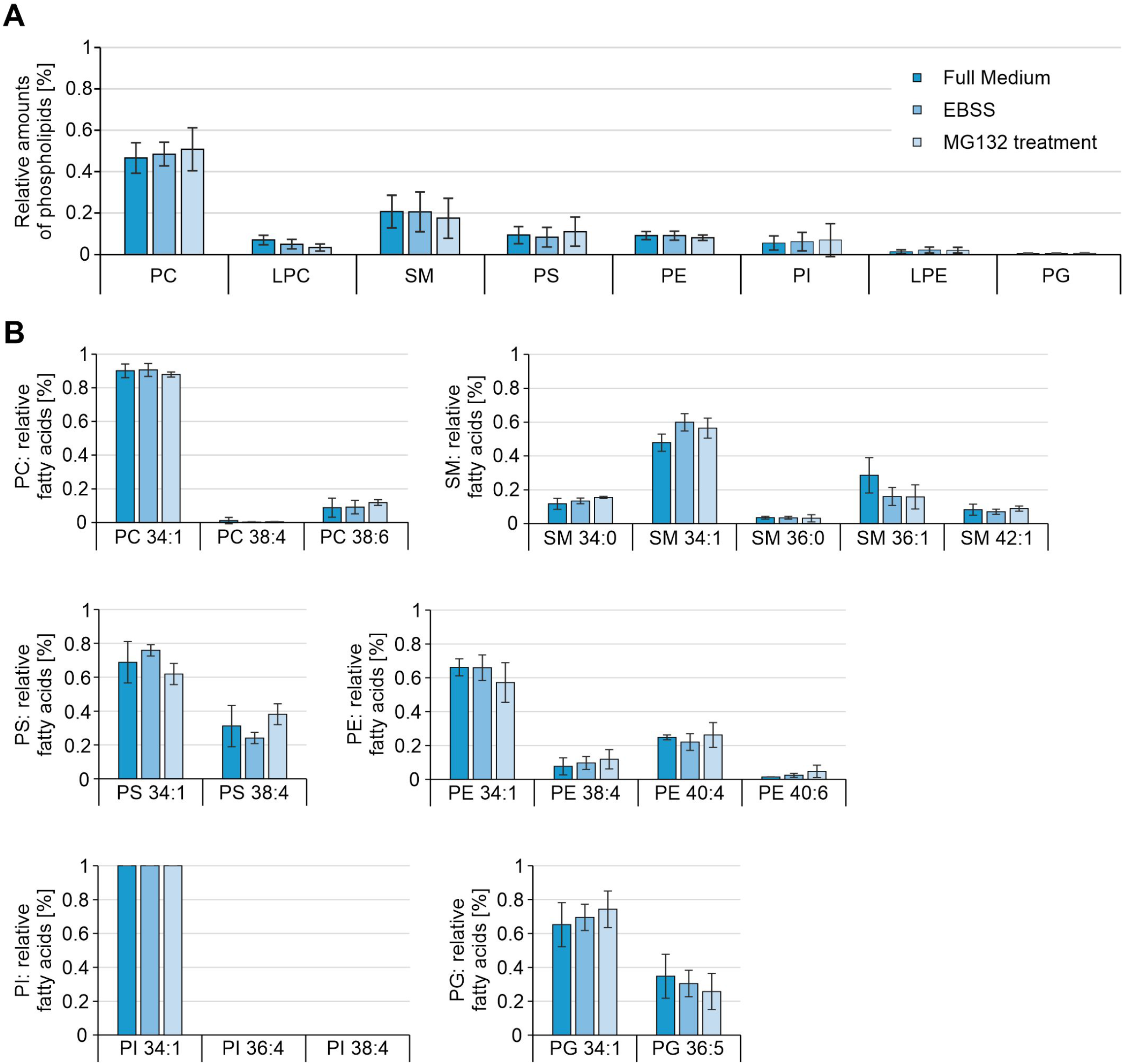
Phospholipid profiles of isolated autophagosomes upon different autophagy conditions. A. Relative phospholipid composition of isolated autophagosomes upon different autophagy conditions. PL ratios were calculated based on their concentration determined in the samples. PC phosphatidylcholine, LPC lysophosphatidylcholine, SM sphingomyelin, PS phosphatidylserine, PE phosphatidylethanolamine, PI phosphatidylinositol, LPE lysophosphatidylethanolamine, PG phosphatidylglycerol. B. Relative fatty acid combinations of identified phospholipids upon different autophagy conditions. Fatty acid ratios were calculated based on their concentration measured in the samples. A+B. Statistics are depicted as mean ± SD; statistical analysis resulted in no significant differences between the different treatments (One-Way ANOVA, n = 3).

Phosphatidylcholine (PC) is the most abundant PL in mammalian cells and mammalian membranes typically show more than 50 % of this particular lipid, which was also reflected by the detected amount of PC in isolated autophagosomes (Figure 4A). The second most frequent PL within the autophagosome fraction was sphingomyelin (SM), the only PL that is not exclusively synthesized within the ER, but is generated from ceramide within the Golgi apparatus. This lipid is commonly found at the Golgi network, the outer leaflet of the plasma membrane, and in the endocytic system (Slotte, 2013). Consistently, besides the ER, the Golgi and endocytic pathways are frequently described to support autophagosome formation (De Tito *et al*, 2020). The detection of SM in autophagic vesicles connects autophagosome membranes to the Golgi apparatus or Golgi-derived vesicles and is in line with a recent publication showing that SM is essential for structure and function of double membrane vesicles responsible for viral replication (Gewaid *et al*, 2020). In addition to their structural role, PLs exhibit also regulatory functions. While present only in lower amounts, PI, PE, and phosphatidylserine (PS) are crucial for efficient autophagosome formation (de la Ballina *et al*, 2020). Phosphorylated forms of PI generated by the PI3-kinase complex, are linked to phagophore generation and elongation (Nascimbeni *et al*., 2017). In an ubiquitin-like conjugation reaction, PE is covalently attached to Atg8 family members and is essential for efficient phagophore maturation and autophagosome function (Noda & Inagaki, 2015). Recent publications showed that lipids are transferred from the ER to the growing phagophore by the activity of ATG2A/B (Osawa et al., 2019; Valverde et al., 2019) and that the presence of negatively charged PS in liposomes accelerates the ATG2B-mediated lipid transfer (Osawa *et al*, 2020).

Concerning fatty acids, the PLs were linked to various combinations. In particular, short chained and (poly-) unsaturated fatty acids were detected at high levels (Figure 4B). Approx. 90 % of PC showed the short fatty acid combination 34:1 that corresponds to the fatty acids 18:1 and 16:0, which was also found in 50 % of SM and in 100 % of PI. Several poly-unsaturated fatty acid variants were also observed. In conclusion, short chained and (poly-) unsaturated fatty acids were most abundant in the PLs present in autophagosomes, which corresponds to recent studies investigating fatty acids in autophagosomes from Drosophila and yeast (Laczko-Dobos *et al*, 2021; Schutter *et al*., 2020).

Autophagosomes are exceptional vesicles with explicit requirements regarding membrane curvature, mobility, and fusion ability. Short chained and (poly-) unsaturated fatty acids increase membrane fluidity and accordingly are supportive for membrane curvature. The PL composition in combination with distinct fatty acid characteristics determine shape and biophysical properties of a membrane and thus autophagosome function is crucially linked to a particular structural and quantitative PL pattern. This is highlighted by the finding that the PL composition is unaffected by the different autophagy conditions, albeit the treatments have significant effects on the rate of autophagosome formation (Appendix Figure S3).

Here, we present a powerful method to isolate native autophagosomes. The FACS-based approach resulted in the purification of intact vesicles, which enabled a profound autophagosome content profiling. We showed that proteasome inhibition modifies autophagic degradation leading to a switch towards a specific selection of cargoes that counteracts the cellular disturbance. Interestingly, nutrient deprivation, while leading to an enhanced autophagic flux, only marginally affects the types of degradation-prone proteins within autophagosomes, emphasizing that the obligation to enhance general protein turnover to recycle building blocks does not require focusing on particular cargoes. Besides protein analysis, isolation of autophagosomes additionally enabled the investigation of phospholipids in autophagosomes. The identified PL pattern was unaffected by different autophagy rates and exhibited a predominant content of short chained, unsaturated fatty acids, highlighting the relevance of a particular PL composition to support the structural and functional requirements of autophagic vesicles.

In summary, we introduce a potent isolation method for native unmanipulated autophagosomes, which paves the way to identify the impact of different autophagy conditions on autophagosome cargo and phospholipid profiles without the requirement of exhaustive cellular fractionation or genetic manipulation.

## Materials and Methods

### Cell culture

HeLa cells were cultivated in DMEM, supplemented with 10 % fetal bovine serum, 1 mM sodium pyruvate, and 1 x antibiotic-antimycotic solution at 37 °C in a 5 % CO_2_-humidified atmosphere. Cell identity was authenticated via STR analysis and cells were regularly tested negative for the presence of mycoplasma using the Venor GeM Mycoplasma PCR detection kit (Minerva Biolabs, 11-1050). Stock solutions of bafilomycin A_1_ (Biozol, TRCB110000) and MG132 (Sigma Aldrich, P0042) were prepared in DMSO. For starvation, cells were treated with serum-free EBSS for 2 h. For proteasome inhibition, cells were incubated with 10 µM MG132 for 8 h. Bafilomycin A_1_ was added to each treatment 2 h before cells were collected.

### Autophagosome isolation

At least 1 × 10^7^ cells were collected using Trypsin/EDTA, centrifuged with 306 x *g* for 4 min and resuspended in PBS supplemented with EDTA-free protease inhibitor (Roche, 11873580001). Cell disruption was performed using a UP50H ultrasonic processor (Hielscher) for 3 x 2 s with an amplitude of 60 %. Lysates were centrifuged with 3.000 x *g* for 10 min at 4 °C, supernatants were collected and centrifuged with 150.000 x *g* (at r_av_) for 30 min at 4 °C using an Optima™ MAX-XP Ultracentrifuge (Beckman Coulter; TLA120.2 rotor). Pellets were resuspended in PBS and 4 µg of primary antibody were added for 30 min, followed by addition of 12 µg Cy5-conjugated secondary antibody for 1 h. The samples were centrifuged with 150.000 x *g* (at r_av_) for 30 min at 4 °C and pellets were resuspended in PBS. Autophagosome sortings were performed using a BD FACSAria III SORP (BD Biosciences) equipped with a 70 µm nozzle and a 1.0 FSC neutral density filter. The autophagosome containing compartment was first established using an FSC/SSC plot on a logarithmic scale, followed by a doublet discrimination gate using SSC-A/W. Autophagosomes were defined as Cy5-positive events (640nm, BP 670/30), whose positivity was conducted according to the background given by an unstained as well as a secondary antibody-only negative control. Cy5-positive autophagosomes were sorted at minimum speed (flow rate < 3.0) maintaining less than 19.000 events per second. Analysis was performed using FlowJo v10.6.1 (BD Biosciences). Primary antibodies: MAP1LC3B (Novus, NB100-2220); GABARAP+GABARAPL1+GABARAPL2 (Abcam, ab109364). Secondary antibody: Cy5 anti-rabbit (ImmunoResearch, 711-175-152).

### Co-localization analysis

Approximately 1 × 10^6^ cells were prepared for FACS analysis as described above using 1 µg primary and 3 µg secondary antibody. Here, a fluorophore-conjugated primary antibody (1 µg) was used in parallel. Autophagosomes were defined as Cy5-positive (640 nm, BP 670/30) as well as Alexa Fluor 488- or Fluorescein-positive events (488 nm, BP 530/30). Positivity was conducted according to the background given by an unstained, a secondary antibody-only, and an unconjugated primary antibody/secondary antibody (Cy5)-only negative control. Analysis was performed using FlowJo v10.6.1 (BD Biosciences). Primary antibodies: MAP1LC3B (Novus, NB100-2220); GABARAP + GABARAPL1 + GABARAPL2 (Abcam, ab109364); MAP1LC3B (fluorescein labeled) (Enzo, BML-PW1205-0025); GABARAP Alexa Fluor 488 (SantaCruz, sc-377300 AF488). Secondary antibody: Cy5 anti-rabbit (ImmunoResearch, 711-175-152).

### Proteinase K digest

5 µg/ml proteinase K were supplemented to purified autophagosomes (5 × 10^6^) and incubated for 15 min on ice. To stop enzyme activity, PMSF (100 mM) was added for 5 min on ice and samples were immediately precipitated using a methanol/chloroform (2:1) precipitation protocol. For positive control, isolated autophagosomes were sonicated using a UP50H ultrasonic processor (Hielscher) for 3 x 3 s with an amplitude of 80 % before proteinase K supplementation. For negative control, 5 µg/ml BSA instead of proteinase K were added to the samples.

### Western blotting

Western blot analyses were performed as previously described (Bekbulat *et al*, 2020). Proteins of isolated autophagosomes were precipitated using a methanol/chloroform (2:1) precipitation protocol and resuspended in urea buffer (8 M urea and 4 % (w/v) CHAPS in 30 mM Tris (pH 8.5 with HCl), including EDTA-free protease inhibitor. Usually 5 µg of total protein or the total protein of 1 to 2 × 10^6^ autophagosomes were subjected to 4 – 12 % NuPage Bis-Tris gels (Thermo Scientific, MP0335) and transferred onto a nitrocellulose membrane using the Trans-Blot Turbo RTA mini nitrocellulose transfer kit (BioRad, 170-4270). After blocking with 5 % fat-free milk in PBS-Tween 20, the membrane was probed with appropriate primary and HRP-conjugated secondary antibodies. Proteins were visualized via chemiluminescence using the Amersham Imager 600 (GE Healthcare Life Science). Quantification was performed using Aida Image Analyzer v4.26 (Raytest). Primary antibodies: MAP1LC3B (Sigma, L7543); GABARAP + GABARAPL1 + GABARAPL2 (Abcam, ab109364); SQSTM1 (Progen, GP62-C); Tubulin α (Sigma, T9026); Actin (Sigma, A5060). Secondary antibodies: Peroxidase anti-guinea pig (ImmunoResearch 706-035-148), Peroxidase anti-mouse (ImmunoResearch 715-035-151), Peroxidase anti-rabbit (ImmunoResearch 711-035-152).

### Immunocytochemistry

Immunocytochemistry was conducted as previously described (Bekbulat *et al*., 2020). Briefly, cells were grown on glass cover slips, fixated using 4 % PFA, and permeabilized with 90 % (v/v) methanol. Unspecific binding sites were blocked with 3 % BSA in PBS followed by incubation with primary antibody, fluorophore-conjugated secondary antibody as well as DAPI. Cells were imaged using the laser scanning microscope LSM710 (Zeiss, Germany). Primary antibody: MAP1LC3B (Nanotools, 0260-100). Secondary antibody: Cy3 anti-mouse (ImmunoResearch, 715-165-151).

### Differential interference contrast microscopy

Isolated autophagosomes were concentrated using Amicon Ultra-15 filters (Merck, UFC901024) and dried on the surface of microscope slides. After sealing the samples with glass cover slips, they were imaged using the widefield microscope AF7000 (Leica).

### Negative stain electron microscopy

5 µl of the FACS sorted autophagosome solution were applied onto freshly prepared Formvar coated copper mesh EM grids and incubated for 1 min at RT. Then, excess liquid was drained by using filter paper and grids were dried at RT. 10 µl of a 2 % (w/v) uranyl acetate (UA) solution was applied to the grid and incubated for 30 sec. The grid was washed 3 x using drops of distilled water and drained using filter paper. Subsequently, the grid was dried over night at RT and analyzed using a TEM900 (Zeiss, Germany) operated at 80 keV and equipped with a wide-angle dual-speed digital 2K CCD camera (TRS Tröndle).

### Sample preparation for LC-MS/MS proteomics

8 × 10^6^ Cy5-positive autophagosomes were denatured with 2 % sodium deoxycholate, 50 mM Tris-HCl pH 8.5, 2.5 mM TCEP, 10 mM chloroacetamide and protease inhibitor cocktail tablet (EDTA-free, Roche) at 95 °C for 10 min. Lysates were prepared with in-StageTip (iST) processing method for LC-MS/MS as previously described by (Kulak *et al*, 2014). Briefly, proteins were digested overnight at 37 °C with 1 vol. of 50mM Tris-HCl pH 8.5 containing LysC (Wako Chemicals) at 1:100 (w/w) ratio and trypsin (Promega V5113) at 1:100 (w/w) ratio. Digestion was stopped with 2 vol. of 1 % TFA in isopropanol. Digested peptides were purified with Empore SDB-RPS (styrenedivinylbenzene - reverse phase sulfonate) disks in stage tip (3M Empore) and were dried for LC-MS/MS.

### Mass spectrometry

Dried peptides of each sample were resuspended in 2 % (v/v) acetonitrile / 1% (v/v) formic acid solution. Peptides were separated with Easy nLC 1200 (ThermoFisher Scientific) using a 30 cm long, 75 μm inner diameter fused-silica column packed with 1.9 μm C18 particles (ReproSil-Pur, Dr. Maisch) and kept at 50 °C using an integrated column oven (Sonation). Individual peptides were eluted by a non-linear gradient from 5 to 40 % B over 120 min, followed by a step-wise increase to 95 % B in 6 min, which was kept for another 9 min and sprayed into a QExactive HF mass spectrometer (ThermoFisher Scientific). Full scan MS spectra (300-1,650 m/z) were acquired with a resolution of 60.000 at m/z 200, maximum injection time of 20 ms and AGC target value of 3 × 10^6^. The 15 most intense precursors were selected for fragmentation (Top 15) and isolated with a quadrupole isolation window of 1.4 Th. MS2 scans were acquired in centroid mode with a resolution of 15,000 at m/z 200, a maximum injection time of 25ms, AGC target value of 1 × 10^5^. Ions were then fragmented using higher energy collisional dissociation (HCD) with a normalized collision energy (NCE) of 27; and the dynamic exclusion was set to 25 s to minimize the acquisition of fragment spectra of already acquired precursors.

### Processing of raw data and statistical analysis

Raw files were analyzed with MaxQuant 1.6.17 with default settings using human trypsin digested “one sequence per gene” proteome database (Homo sapiens SwissProt database [TaxID:9606, version 2020-03-12]) with label free quantification (Cox & Mann, 2008). Carbamidomethyl on cysteines were set as fixed modification and acetyl on the protein N-term and methionine oxidation on methionines were set as variable modifications. Protein quantification and data normalization relied on the MaxLFQ algorithm implemented in MaxQuant and used for statistical analysis (Cox *et al*, 2014). Proteins only identified by a single modified peptide or matching to the reversed or contaminants databases were removed. Only well-quantified proteins and showing no missing value in any of the samples were used for statistical analysis. Significantly altered proteins were determined by a two-sided, unpaired Student’s *t*-test (p-value < 0.05) adding minimum fold-change cut-off (> = 0.5) with GraphPad Prism 6 or Microsoft Excel 2016. No fold change could be calculated are indicated as not determinable (nd) in the volcano plots.

### Network analysis

Protein pathway network analysis was performed with Cytoscape (Shannon *et al*, 2003) (version 3.8.2) in combination with StringApp, which is using STRING database (Doncheva *et al*, 2019). KEGG databases were used to obtain pathway enrichment (Kanehisa *et al*, 2010).

### Data availability

The mass spectrometry proteomics data have been deposited to the ProteomeXchange Consortium via the PRIDE partner repository (Perez-Riverol *et al*, 2019) with the dataset identifier PXD024419 (Username; Password: 18d5GdUA).

### Lipidomics

Lipids were extracted from autophagosomes using a liquid-liquid extraction method previously described (Lerner *et al*, 2017). Briefly, autophagosome preparations were allowed to thaw at 4 °C. 1000 µl methyl tert buthyl ether (MTBE) /methanol (10:3, v/v) spiked 10 µl of internal standard mixture for PLs and SMs and 250 µl 0.1 % formic acid containing 5 µm THL/URB and 10 µg/ml BHT were added and samples were subjected to homogenization for 20 sec at 6000 rpm with a Precellys 24 (Bertin Technologies, Montigny-le-Bretonneux, France), were vortexed for 30 sec at 4 °C, and centrifuged for 10 min at 13000 rpm at 4 °C. Then samples were allowed to freeze at -20 °C for 15 min and the upper organic phase containing lipids was recovered and dried under a gentle stream of nitrogen at 37 °C. The dried lipid extracts were re-solubilized in 90 µl methanol and stored at -20 °C until analysis. The lower, aqueous phase containing autophagosomal proteins was stored at -20°C for further BCA assay protein content determination, which was used for normalization of PL values. Lipids were analyzed by liquid-chromatography multiple reaction monitoring (LC/MRM) on a QTRAP 55500 mass spectrometer (AB Sciex, Darmstadt, Germany) operated in positive negative ion mode switching. The chromatographic, ionization and detection conditions were set as previously described (Lerner *et al*., 2017). The MRM transitions for the analysis of selected PLs and SMs as well as calibration standards used for quantification were set as previously reported (Lerner *et al*, 2019; Lerner *et al*., 2017). For LC/MRM analysis, 3 µl water were added to a 27 µl methanolic solution of extracted lipids and 10 µl of this solution were injected into the LC/MRM instrument. Lipids were quantified using Multiquant 3.0.3 software and determined lipid levels were normalized to the protein content of the autophagosomes.

### Statistics

In dependence of normal distribution or variance differences, statistical significance between the samples were determined by unpaired Student’s *t*-test or One-Way ANOVA using SIGMA STAT (SPSS Science). The results are shown as mean ± standard deviation (SD). Statistical significance was accepted at a level of p < 0.05 (p value ≤ 0.05 = *, ≤ 0.01 = **, ≤ 0.001 = ***).

## Acknowledgments

We would like to thank Drs Stefanie Möckel and Jesús Gil-Pulido of the Flow Cytometry Core facility as well as Drs Sandra Ritz and Petri Turunen of the Microscopy and Histology Core facility, both Institute of Molecular Biology, Mainz, for scientific help and discussions. In addition, we thank Marion Basoglu for help with EM experiments conducted in the FB15 EM Facility of the Goethe University Frankfurt, which is part of the Frankfurt Center for Electron Microscopy (FCEM). This work was funded by the Deutsche Forschungsgemeinschaft (DFG, German Research Foundation) – Project-ID 259130777-SFB1177. Lipidomic experiments were financially supported by the Core Facility of the Clinical Lipidomics Unit.

## Author contributions

D.S. was supported by P.H.D. and H.H. and performed all the experiments, except TEM, conducted by S.E., proteomics, conducted by S.B., G.T., and C.M., and lipidomics, conducted by L.B. All authors participated in data analysis

and discussion. D.S., C.B., and A.K. conceived the study and wrote the manuscript.

## Conflict of interest

The authors declare no conflict of interest.

## Figure legends

**Appendix Figure S1.**
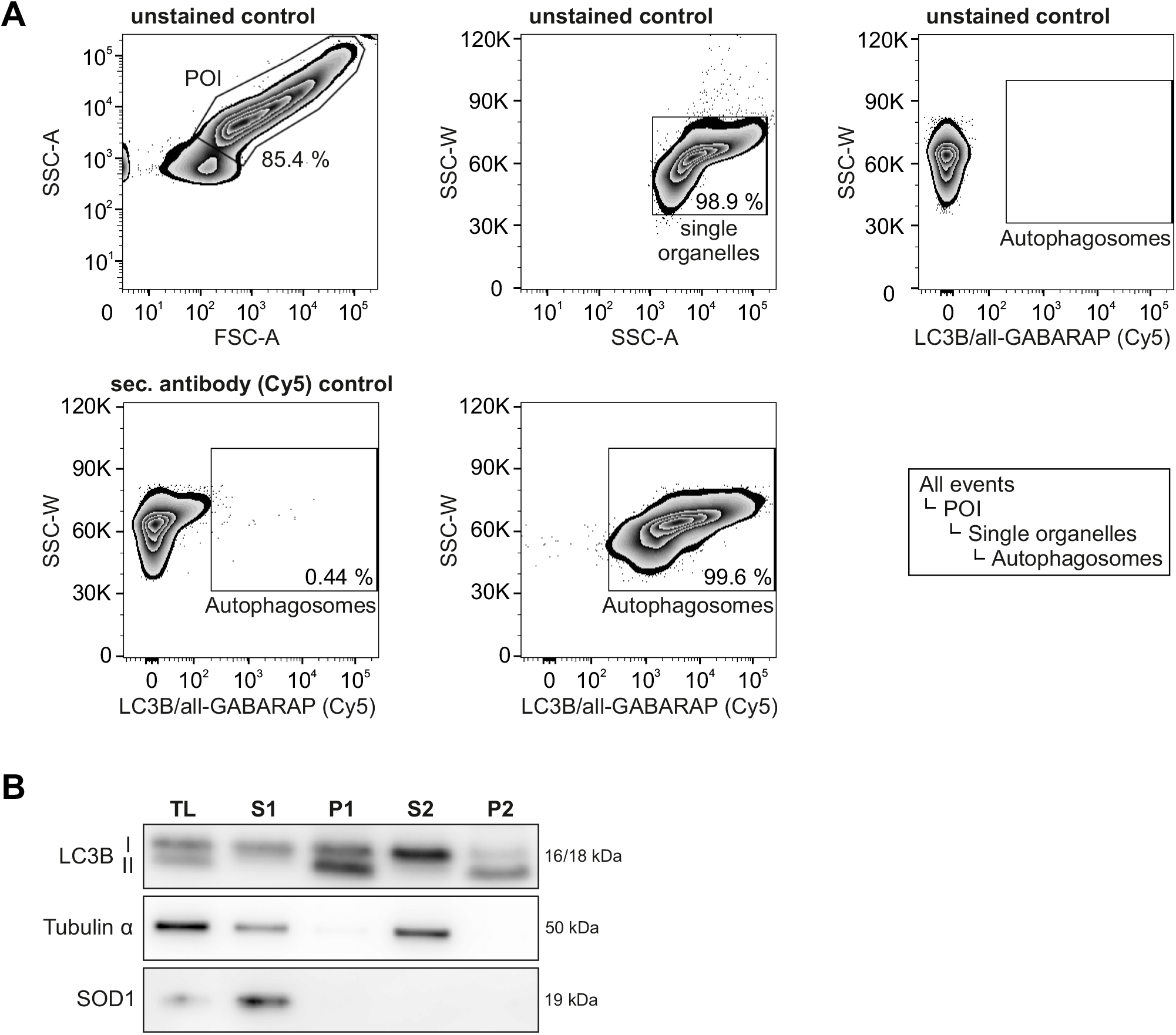
FACS-based autophagosome isolation. A. Representative images of the FACS gating strategy. The population of interest (POI) containing autophagosomes was first established using a FSC/SSC plot on a logarithmic scale, followed by a doublet discrimination gate using SSC-A/W. Cy5-positive events (640nm, BP 670/30) were defined as autophagosomes, whose positivity was conducted according to the background given by an unstained as well as a secondary antibody-only negative control. B. Western blot analysis of different steps of the isolation protocol. TL total lysate, S supernatant, P pellet. Shown is a representative blot of 3 independent approaches.

**Appendix Figure S2.**
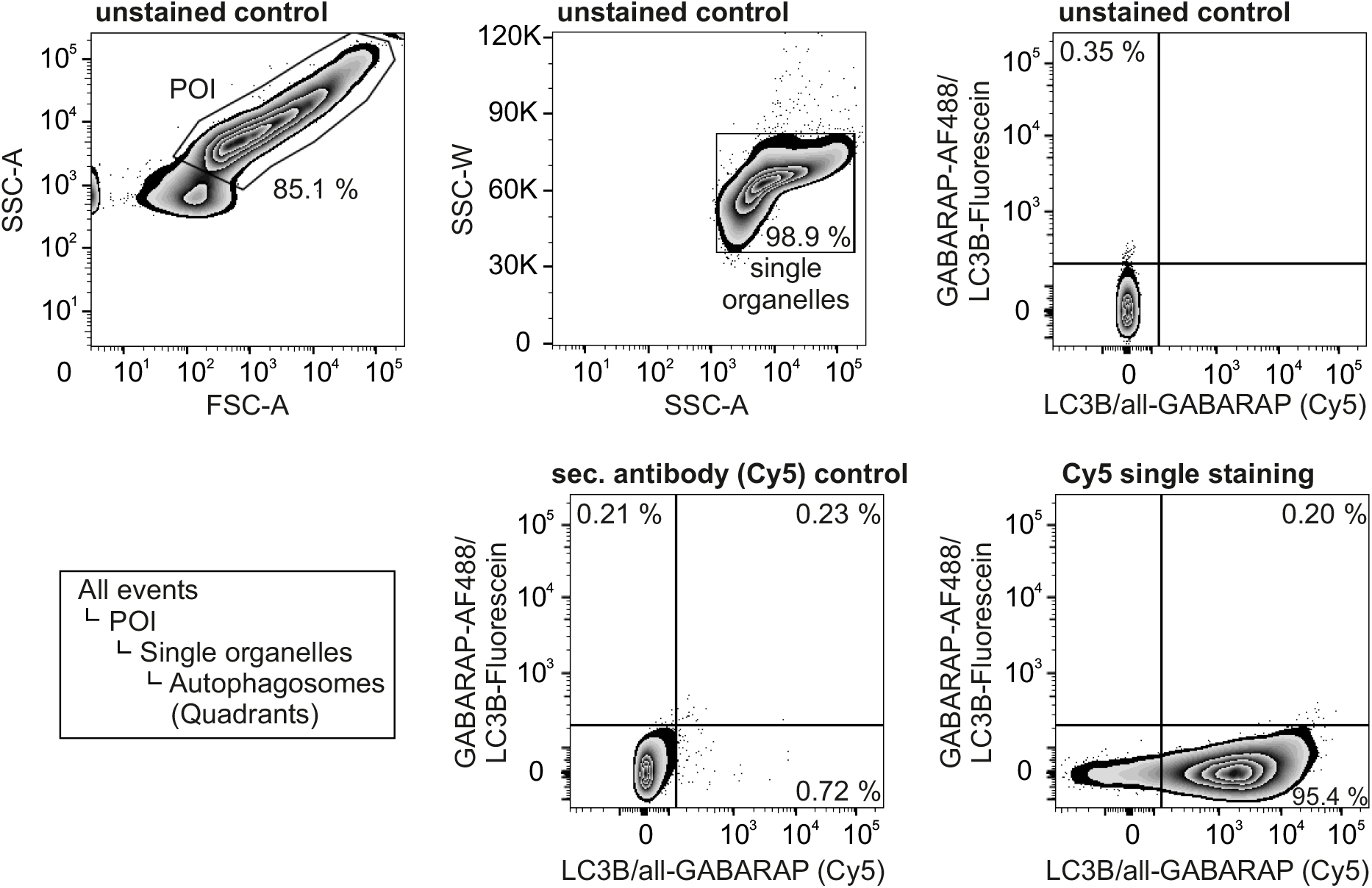
Gating strategy of the co-localization assay. The population of interest (POI) containing autophagosomes was first established using a FSC/SSC plot on a logarithmic scale, followed by a doublet discrimination gate using SSC-A/W. Autophagosomes were defined as Cy5-positive events (640 nm, BP 670/30) and Alexa Fluor 488- or Fluorescein-positive events (488 nm, BP 530/30). Their positivity was conducted according to the background given by an unstained, a secondary antibody (Cy5)-only as well as a primary antibody/secondary antibody (Cy5)-only negative control.

**Appendix Figure S3.**
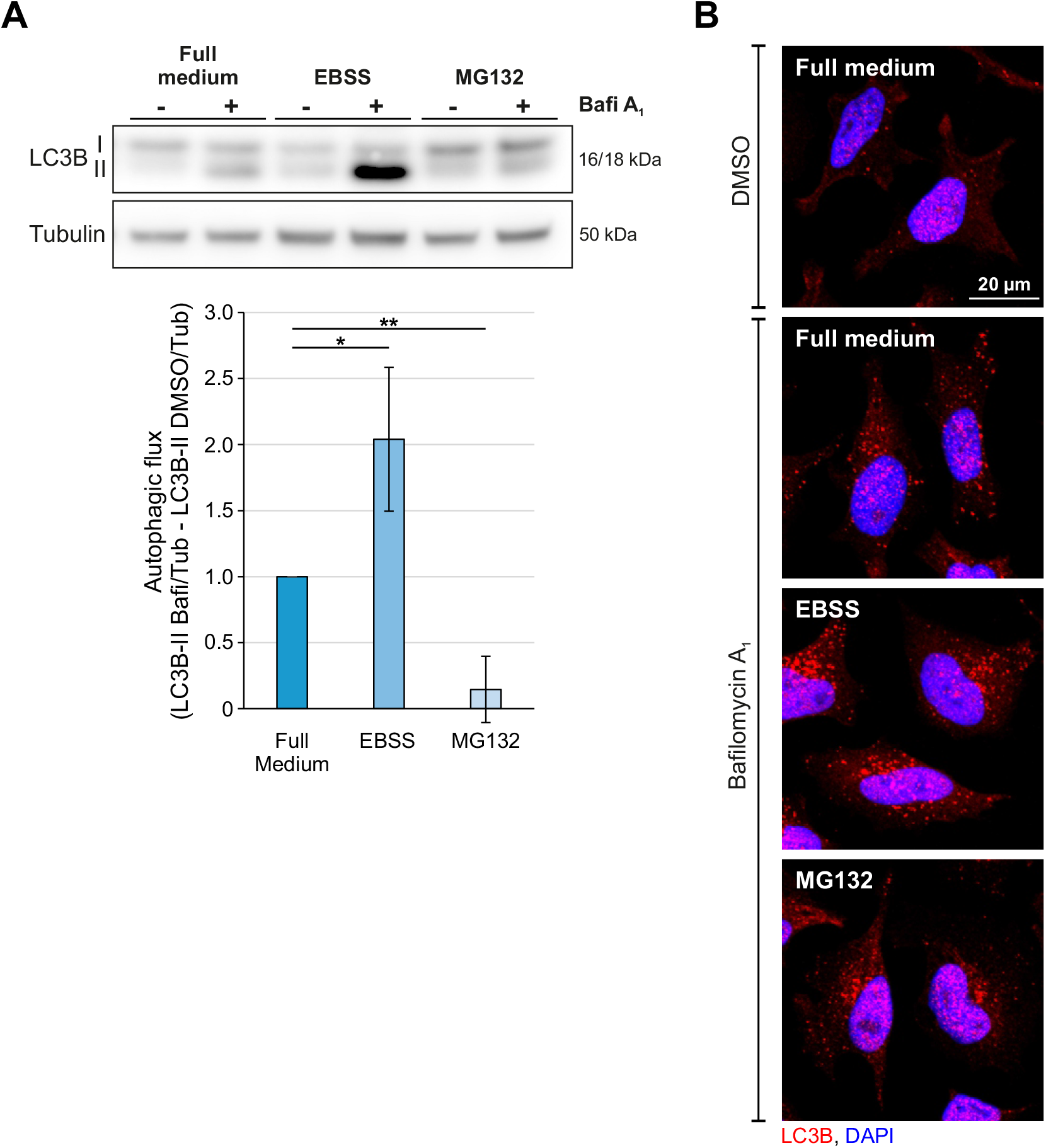
Effects of different treatments on autophagy. A. Western blot analysis of autophagic activity (LC3B-II flux) upon different treatment paradigms. Cells were treated with DMSO or bafilomycin A1. LC3B-II levels were corrected over the loading control tubulin. Statistics are depicted as mean ± SD. (n = 3, *p ≤ 0.05; **p ≤ 0.01). B. Representative confocal images of immunocytochemical stainings of LC3B (red) upon different treatment paradigms. Nuclei were stained with DAPI (blue).

**Appendix Figure S4.**
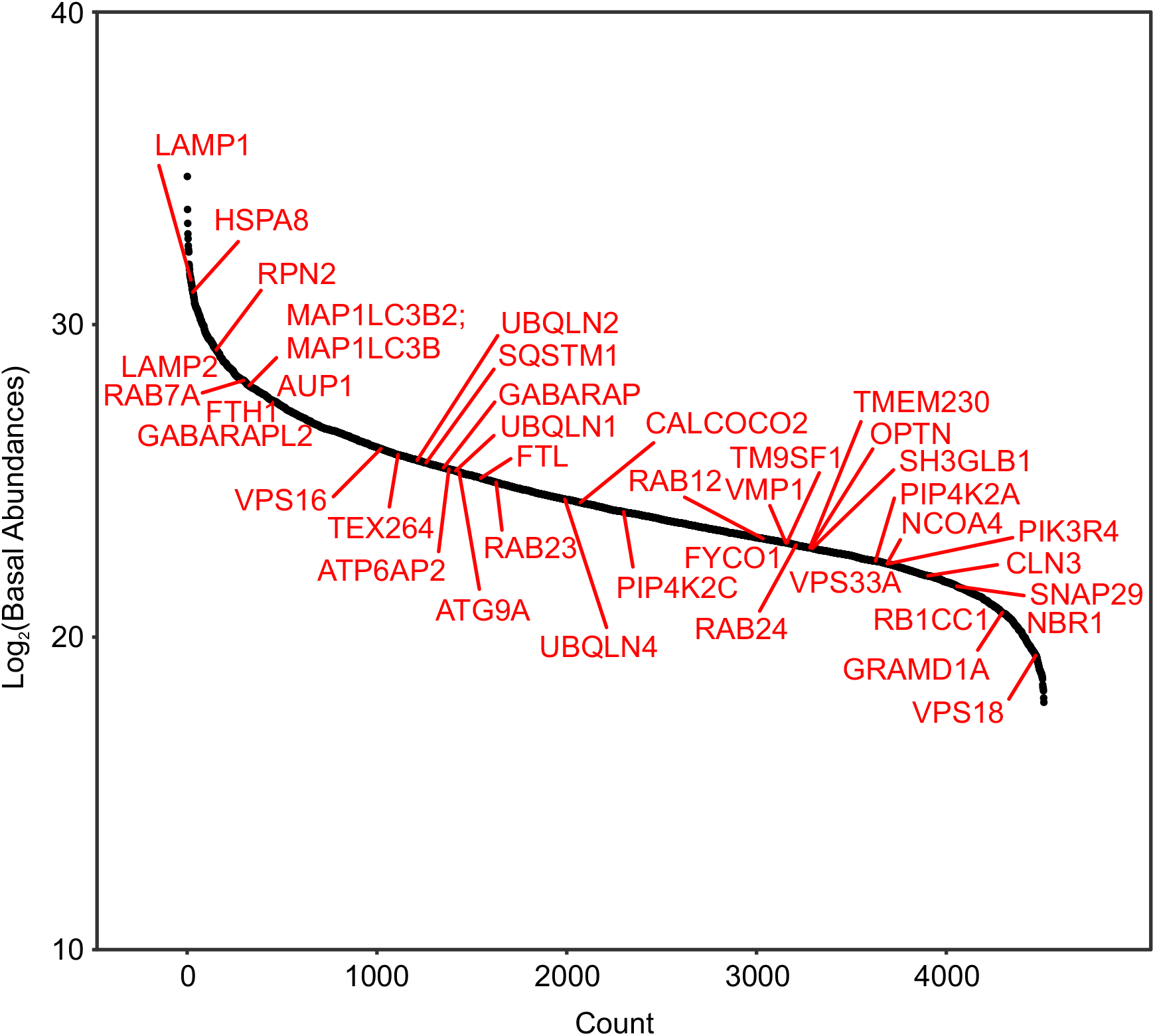
Autophagosomal proteins in basal autophagosomes. Abundance plot of autophagosome proteins detected in isolated autophagosomes upon basal conditions.

**Appendix Figure S5.**
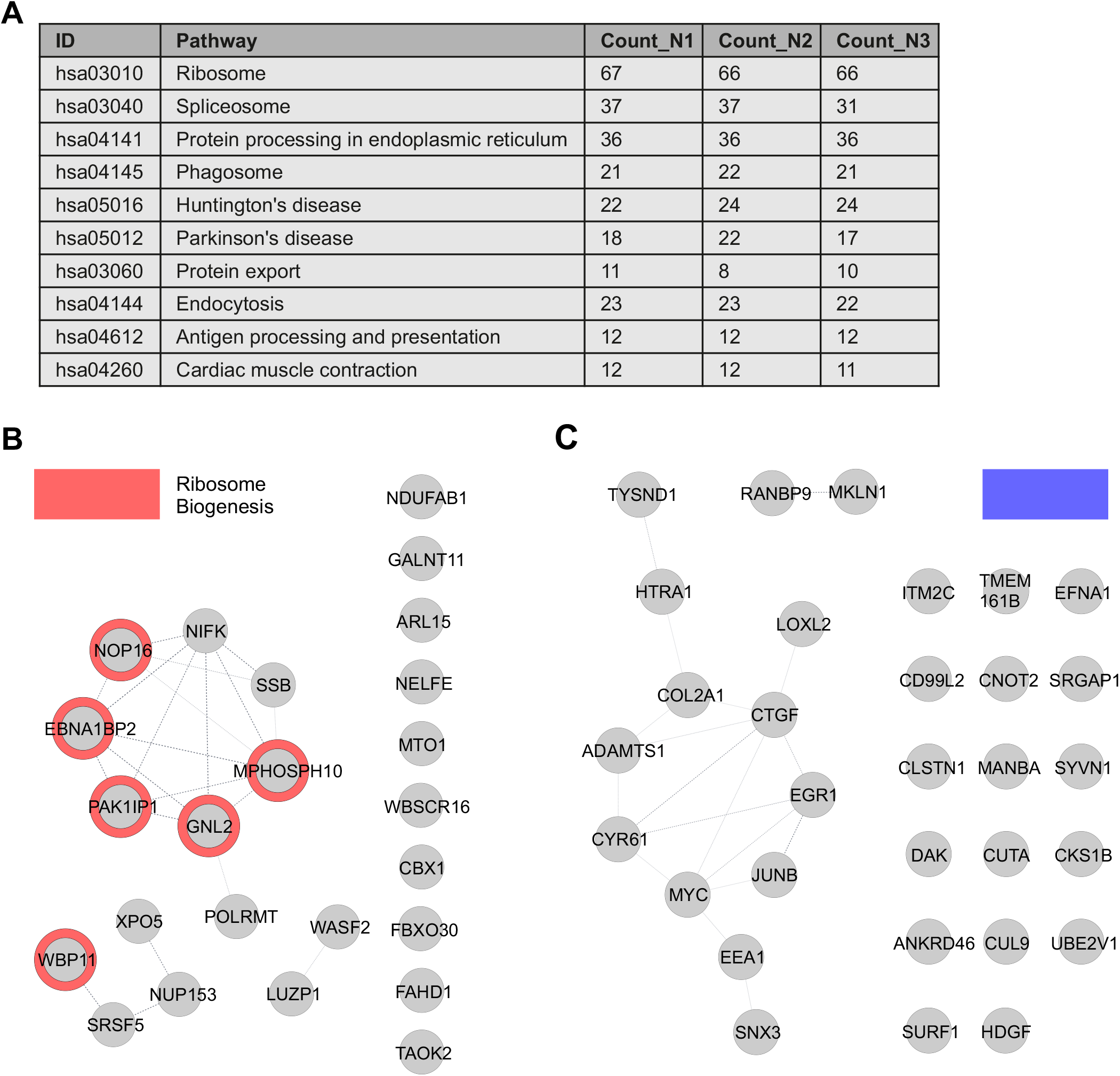
Cargo analysis of autophagosomes upon basal and starved conditions. A. KEGG pathways are presented with the number of proteins found in the data set of each autophagosome analysis from 3 independent autophagosome isolations upon basal autophagy conditions. B. String analysis of proteins with enhanced abundance in autophagosomes upon EBSS treatment. C. String analysis of proteins with reduced abundance in autophagosomes upon EBSS treatment.

**Appendix Figure S6.**
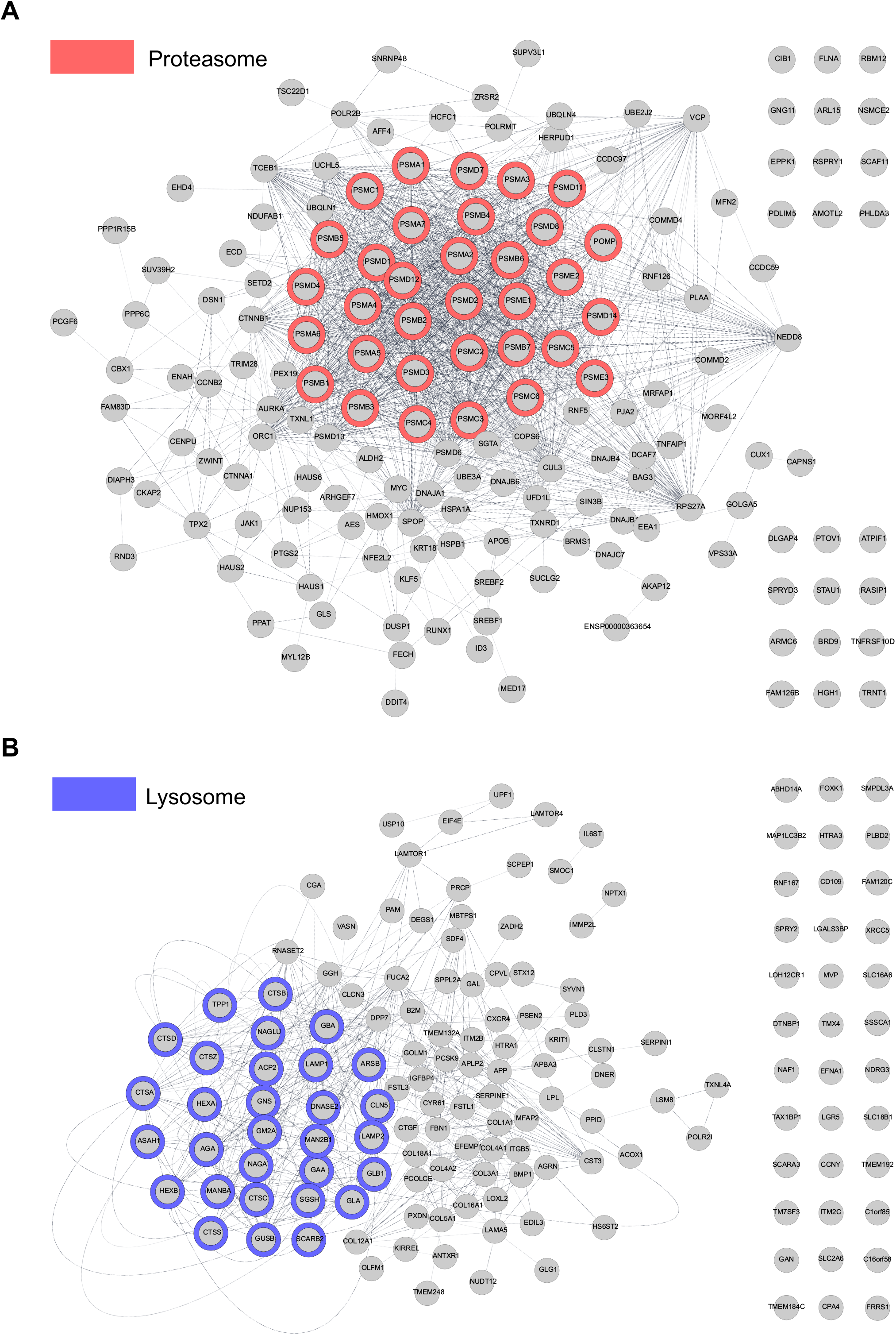
Cargo analysis of autophagosomes upon proteasome inhibition. A. String analysis of proteins with enhanced abundance in autophagosomes upon MG132 treatment. B. String analysis of proteins with reduced abundance in autophagosomes upon MG132 treatment.

**Appendix Figure S7.**
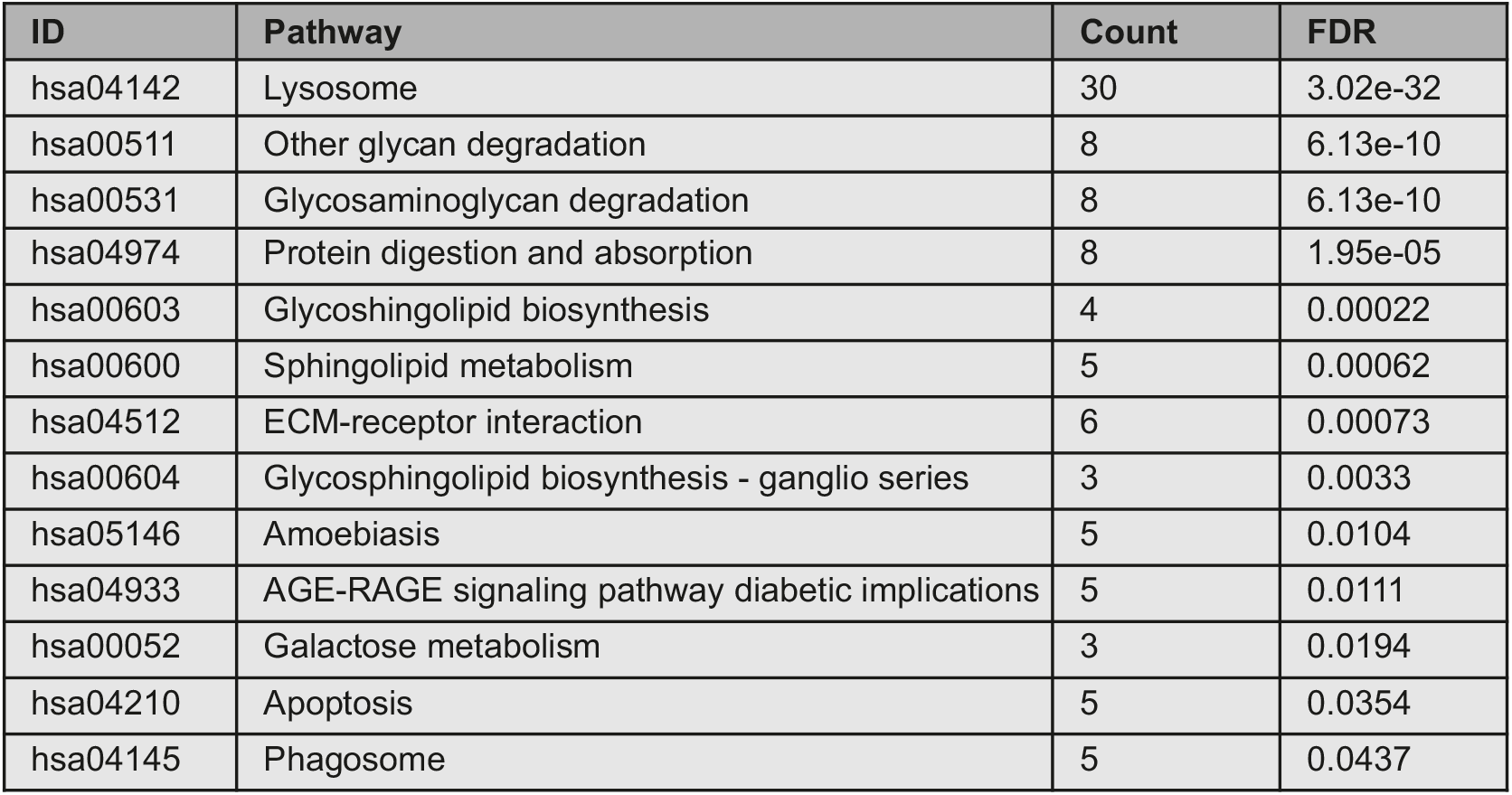
Pathways analysis of proteins with reduced appearance in autophagosomes upon MG132 treatment. KEGG pathway analysis of proteins with reduced autophagosome levels upon MG132 treatment. Pathways are presented with the number of proteins found in in the data set and computed FDRs for enrichment.

